# Genomic diversity and evolution in the Hawaiian Islands endemic Kokia (Malvaceae)

**DOI:** 10.1101/2024.04.17.589907

**Authors:** Ehsan Kayal, Mark A. Arick, Chuan-yu Hsu, Adam Thrash, Mitsuko Yorkston, Clifford W. Morden, Jonathan F. Wendel, Daniel G. Peterson, Corrinne E. Grover

**Affiliations:** Ecology, Evolution, and Organismal Biology Dept., Iowa State University, Ames, IA, 50011, USA; Institute for Genomics, Biocomputing & Biotechnology, Mississippi State University, Mississippi State, MS 39759, USA; School of Life Sciences, University of Hawai‘i, Honolulu, HI 96822, USA

**Keywords:** Kokia cookei, K. drynarioides, K. kauaiensis, Hawaiian forest conservation

## Abstract

Island species are highly vulnerable due to habitat destruction and their often small population sizes with reduced genetic diversity. The Hawaiian Islands constitute the most isolated archipelago on the planet, harboring many endemic species. *Kokia* is an endangered flowering plant genus endemic to these islands, encompassing three extant and one extinct species. Recent studies provided evidence of unexpected genetic diversity within *Kokia*. Here, we provide high quality genome assemblies for all three extant *Kokia* species, including an improved genome for *K. drynarioide*s. All three *Kokia* genomes contain 12 chromosomes exhibiting high synteny within and between *Kokia* and the sister taxon *Gossypioides kirkii*. Gene content analysis revealed a net loss of genes in *K. cookei* compared to other species, whereas the gene complement in *K. drynarioides* remains stable and that of *K. kauaiensis* displays a net gain. A dated phylogeny estimates the divergence time from the last common ancestor for the three *Kokia* species at ∼1.2 million years ago (mya), with the sister taxa [*K. cookei + K. drynarioides*] diverging ∼0.8 mya. *Kokia* appears to have followed a stepping-stone pattern of colonization and diversification of the Hawaiian Archipelago, likely starting on low or now submerged older islands. The genetic resources provided may benefit conservation efforts of this endangered endemic genus.

## Introduction

Human-driven biodiversity loss has greatly contributed to the decline in many species, leading to a rate of species loss reminiscent of past mass extinction events (Storch *et al*. 2022). Species occupying island habitats with small population sizes are particularly vulnerable to sharp reductions in genetic diversity and inbreeding depression (Cowie *et al*. 2022). One such island habitat is the Hawaiian archipelago, whose distance of ∼4,000 km from the nearest continent makes it the most isolated major island chain in the world. Many species endemic to these islands are threatened by extinction through a combination of habitat destruction, invasive species, and predation (Chynoweth *et al*. 2010; Hibit and Daehler 2019). One such example is the genus *Kokia* (Figure 1), an endangered genus of small trees endemic to the Hawaiian Islands and belonging to the Malvaceous tribe Gossypieae, which also contains the economically important cotton genus (Seelanan *et al*. 1997; Hu *et al*. 2021). Once a prevalent part of the xeric-mesic forests of the Hawaiian Islands, *Kokia* has experienced significant declines in diversity: *Kokia drynarioides*, originally described in the dry forests and lava fields of Hawai‘i Island; *K. kauaiensis*, found in the mesic forest of western Kaua‘i Island; *K. cookei* endemic to the Western end of the Moloka‘i Island and now existing only as graft on *K. drynarioides*; and the extinct *K. lanceolata* (Sherwood and Morden 2014).

**Figure 1:**
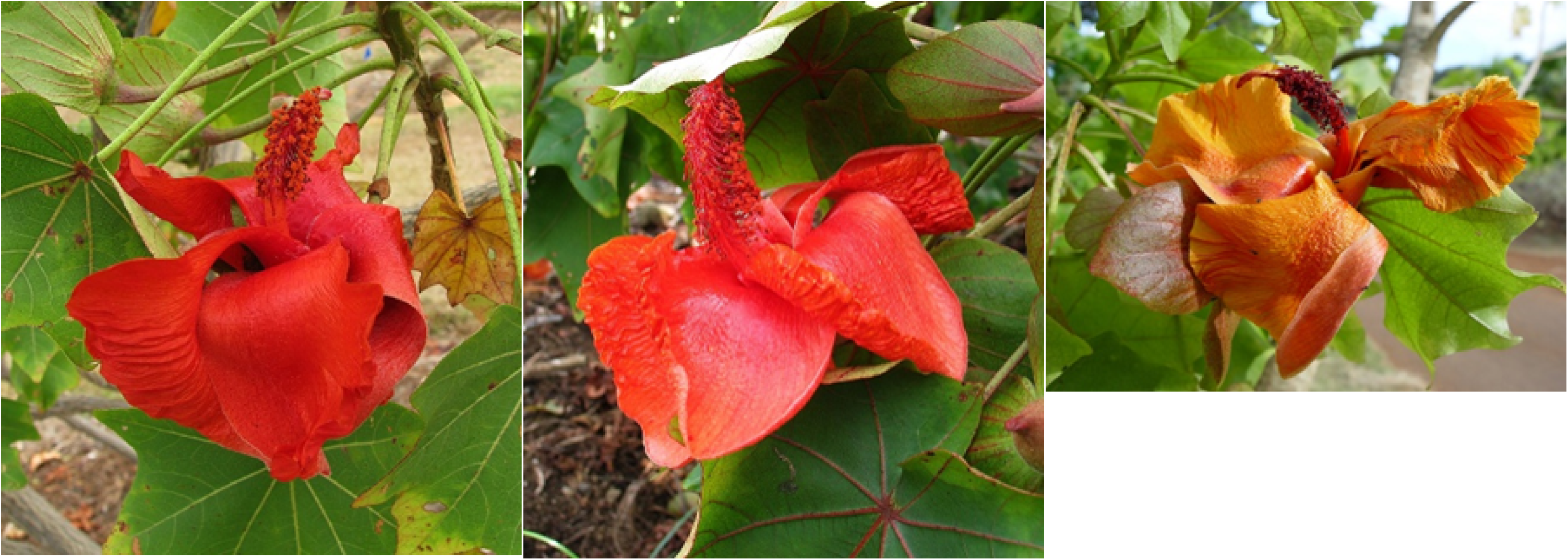
Flowers of the three *Kokia* species sequenced and presented in this study. A) *K. cookei*; origin: Moloka‘i Island; status: extinct in the wild, all individuals believed to be derived from a single plant, one of the most endangered plant species. Photo by David Eichoff (<u>CC BY</u> <u>2.0 DEED</u>). B) *K. drynarioides*; origin: Hawai‘i Island; status: critically endangered. Photo by David Eichoff (<u>CC BY 2.0 DEED</u>). C) *K. kauaiensis*; origin: Kaua‘i Island; status: critically endangered, only 45-50 individuals left in the wild. Photo from National Tropical Botanical Garden, https://ntbg.org/.

Despite its historical importance to the Hawaiian Islands, few studies have focused on genetic diversity of *Kokia* within a conservation framework, using only randomly amplified polymorphic DNA (RAPD) and/or a small number of genetic markers (Sherwood and Morden 2014; Morden and Yorkston 2018). These studies found a higher than expected genetic diversity in *K. kauaiensis* and a surprising population structure that does not match the geography of the islands (Sherwood and Morden 2014). Furthermore, these studies found some level of genetic diversity among *K. cookei* individuals, despite their extreme genetic bottleneck as propagated clones derived from a single initial grafted individual. Previous phylogenetic analyses support the stepping-stone dispersal model, which suggests that *Kokia* spread across the Hawaiian Islands as new islands emerged (Morden and Yorkston 2018). While these previous studies provide valuable insight into the evolution and current status of *Kokia* species, our present understanding is limited by the anonymous and low-throughput nature of the genetic data previously available.

Recently, the first *Kokia* genome was obtained from a *K. drynarioide*s specimen maintained in the Iowa State University greenhouse (Grover *et al*. 2017; Udall *et al*. 2019). Analysis of this genome sequence in conjunction with the closely related *Gossypioides kirkii* genome (∼ 5 million years divergence; (Grover *et al*. 2017)) revealed remarkable divergence between these two genera, particularly in the genic fraction, which exhibited only ∼70 % overlap in gene content. Here we extend this analysis to evaluate the divergence among *Kokia* species using two new high-quality genome assemblies for the other two extant *Kokia* species (i.e., *K. cookei* and *K. kauaiensis*), as well as an improved assembly for *K. drynarioide*s. We reevaluate genomic diversity in *Kokia* and provide foundational resources that are relevant to conservation efforts in this endangered genus.

## Methods & Materials

### Plant material and sequencing methods

#### DNA extraction and sequencing

Fresh leaf tissue was harvested from *K. cookei* (WAI 16c69) and *K. kauaiensis* (WAI 19s9) growing at Waimea Valley (arboretum and botanical garden in Haleiwa, HI, USA) and transported to the mainland under permit I2665. Fresh leaf tissue was also collected from the *K. drynarioides* growing in the Iowa State University greenhouse. All tissue was shipped on ice to Mississippi State University for DNA extraction and sequencing.

The high molecular weight (HMW) nuclear genomic DNA from each species was extracted using modified nuclear genomic DNA isolation procedure combining the nuclei isolation method described in Paterson *et al*. (1993) and the genomic DNA extraction method using Qiagen Plant DNeasy Mini kit (Qiagen, Germantown, MD, USA) following the manufacturer’s instruction with minor modification (Paterson *et al*. 1993). Briefly, 200 mg of leaf tissues were ground into fine powders with a mortar and pestle in liquid nitrogen. The tissue powders were suspended with 1.5 ml of ice-cold extraction buffer and transferred into a microcentrifuge tube (Paterson *et al*. 1993). The nuclei were pelleted by centrifuging at 4°C with the speed of 2,700x g for 20 minutes. After removing the supernatant, the pelleted nuclei were suspended in 400 µl of AP1 buffer (from Qiagen Plant DNeasy Mini Kit) with 4 µl of RNase A (100 mg/ml) (Qiagen, Germantown, MD, USA), The extraction procedure was then followed by the manufacturer’s manual. To get high molecular weight genomic DNA, the centrifugation speed was decreased from 6,000x g to 4,500x g after applying the cell lysate into the DNeasy Mini spin column, followed by elution of the DNA with 50 µl of 10 mM Tris-HCl buffer (pH 8.5). The concentration and purity of extracted nuclear genomic DNA was measured by the NanoDrop One spectrophotometer (Thermofisher Scientific, Waltham, MA, USA) and Qubit fluorometer with the Qubit dsDNA BR assay kit (Life Technologies, Grand Island, NY). The quality of nuclear genomic DNA was validated by agarose gel electrophoresis.

The nuclear genomic DNA from *K. drynarioides* was fragmented with g-Tube (Covaris, Woburn, MA, USA) by centrifuging at 2,300x g for 1 min to generate the mean fragment size of 13-14 kb. The fragmented DNA was subjected to Nanopore DNA library prep using a Genomic DNA Ligation Sequencing Kit (SQK-LSK109; Oxford Nanopore Technologies, Oxford, UK) based on the manufacturer’s protocol and followed by sequencing on a Nanopore GridION sequencer using the MinION R9.4.1 flow cell (Oxford Nanopore Technologies, Oxford, UK) for 48-hr run. The raw sequencing data produced from five MinION flow cells were used for the whole genome assembly.

For both *K. cookei* and *K. kauaiensis*, three library size selection protocols were used to generate different size ranges of DNA fragments, including the mean fragment size of 13-15 kb, the fragment size range from 10 to 46 kb, and the fragment size range from 10 to 150 kb. In brief, the g-Tube (Covaris, Woburn, MA, USA) was used to shear 3 µg of nuclear genomic DNA by centrifuging at 2,000x g for 1 min to get DNA fragments with mean size of 13 to 15 kb, or to select the size range from 10 to 46 kb and 10 to 150 kb using SageELF size fractionater (Sage Science, Beverly, MA, USA) and BluePippin size selection system(Sage Science, Beverly, MA, USA), respectively. The Nanopore DNA libraries were prepared from the fragmented DNAs by using a Genomic DNA Ligation Sequencing Kit (SQK-LSK112; Oxford Nanopore Technologies, Oxford, UK) and sequenced on a Nanopore GridION sequencer using the MinION R10.4.1 flow cell (Oxford Nanopore Technologies, Oxford, UK) for 72-hr run based on the manufacturer’s protocol. The raw sequencing data produced from six MinION flow cells (two for each size selection method) per species were used for the whole genome assembly.

For Hi-C sequencing, five hundred mg of *Kokia* leaf tissues (for both *K. cookei* and *K. kauaiensis*) were ground into powders in liquid nitrogen using a mortar and pestle and directly used for constructing the Hi-C library using the Proximo Hi-C Plant Kit (Phase Genomics, Seattle, WA, USA) followed by the manufacturer’s procedure. The quantity and quality of library were validated by using the Qubit fluorometer with the Qubit dsDNA HS assay kit (Life Technologies, Grand Island, NY) and Agilent Bioanalyzer 2100 with Agilent DNA 1000 Kit (Agilent Technologies, Santa Clara, CA), respectively. The Hi-C library samples were shipped to Novogene Corporation (Novogene Inc., Sacramento, CA, USA; https://www.novogene.com/us-en/) for sequencing with one lane of Pair-End 150 (PE150) sequencing run per species using Illumina HiSeq X-Ten sequencer (Illumina, Sand Diego, CA, USA).

### Genome Assembly

Raw nanopore data for *K. drynarioides* were base called via the Oxford Nanopore Technology (ONT) guppy basecaller v.6.4.6 using the super accuracy plant model (dna-r9.4.1_450bps_sup_plant) and then filtered for lambda control sequences and sequences shorter than 4 kbp using devour (https://gitlab.com/IGBB/devour). The filtered reads were assembled into contigs using canu v.2.1 (Koren *et al*. 2017). Since canu performs correction during the assembly process, additional polishing was not run.

The nanopore data for *K. kauaiensis* and *K. cookei* were base-called with guppy using the super accuracy model (dna_r10.4.1_e8.2_260bps_sup). Contigs were assembled for each species using hifiasm v0.18.5-r499 (Cheng *et al*. 2021, 2022) in conjunction with the base-called reads and Hi-C library for each species. Primary contigs for each species were corrected using medaka v1.7.1 (https://github.com/nanoporetech/medaka) with the corresponding base-called reads.

Each assembly was scaffolded using yahs v.1.1 (Zhou *et al*. 2023) with their respective raw Hi-C libraries that had been aligned to the contigs with bwa v.0.1.17 (Li 2013), deduplicated with samblaster v.0.1.29 (Faust and Hall 2014), and sorted with samtools v.1.17 (Petr Danecek, James K Bonfield, Jennifer Liddle, John Marshall, Valeriu Ohan, Martin O Pollard, Andrew Whitwham, Thomas Keane, Shane A McCarthy, Robert M Davies, Heng Li 2021). Scaffolds were aligned to the *G. kirkii* reference (downloaded from cottongen.org; (Udall *et al*. 2019)) using minimap2 v.2.17 (Li 2018, 2021) and visualized with dotplotly (https://github.com/tpoorten/dotPlotly). Scaffolds that spanned chromosomes were split using agptools v.0.0.1 (https://github.com/WarrenLab/agptools).

The corrected yahs-scaffolded assemblies were further scaffolded into chromosomes via ragtag v.2.1.0 (Alonge *et al*. 2022) using the minimap2 aligned scaffolds to the *G. kirkii* reference. The agp files produced by yahs and ragtag were merged for each species, producing contig linkages for each chromosome. These linkages were manually evaluated and adjusted based on the contact maps plotted with hic-viz (https://github.com/IGBB/hic-viz) using the contig aligned Hi-C library used for yahs scaffolding. To aid in the manual adjustment, magpie (https://github.com/IGBB/magpie) was developed after finishing the assembly for *K. drynarioides*. The final assemblies were produced from the contigs and the linkages using agptools.

The complete code, scripts, and parameters used for assembly can be found at https://github.com/IGBB/Kokia/tree/master/wgs/1-assembly.

### Repeat and gene annotation

RepeatModeler v.2.0.5 (Flynn *et al*. 2020) was used to create a species-specific repeat database for each of the three *Kokia* assemblies and the *G. kirkii* reference. RepeatMasker v.4.1.5 (https://www.repeatmasker.org/) annotated and masked the repeats in each genome. Genes were predicted for the masked genomes using BRAKER3 v.1.0.4.1 (Gabriel *et al*. 2023) with the OrthoDB v.11 Viridiplantae protein database (Kuznetsov *et al*. 2023). InterproScan v.5.65-97.0 (Jones *et al*. 2014) was used to functionally annotate all predicted peptide sequences for each genome. The complete code, including specific parameters, can be found at https://github.com/IGBB/Kokia/tree/master/wgs/2-annotation

### Comparisons among extant Kokia species

We used GENESPACE v.1.2.3 (Lovell *et al*. 2022) to compare the newly produced *Kokia* genomes using *Gossypioides kirkii* as an outgroup. To do so, we limited our analyses to sequences and genes assembled into chromosomes. GENESPACE was run with default parameters and was restricted to sequences and annotations that assembled into the twelve chromosome, removing data that fell into non-chromosome scaffolds, producing a list of syntenic orthologs (SynOGs). The *plot_riparian* module was used to create a genomic map of the twelve chromosomes. Copy number variation (CNV) was evaluated using the “pangenome” outputs from GENESPACE. Runs with *Kokia* only and *Kokia* + *Gossypioides* input taxa were both used to investigate CNV within *Kokia* and between *Kokia* and *Gossypioides*, respectively. In each case, the number of genes per species for each identified SynOG was counted and reported on the species phylogeny. GENESPACE also created a list of single-copy orthogroups (SCOGs) produced by the OrthoFinder module that were used for the rest of the analyses.

To reconstruct the phylogenetic relationships between the *Kokia* species, we amino acid sequences were aligned for individual SCOGs using mafft v.7.508 (--reorder --auto) (Katoh and Standley 2013); amino acids alignments were used to generate single-gene nucleotide alignments with Pal2Nal v.14.1 (Suyama *et al*. 2006); each nucleotide alignment was filtered with gblocks v.0.91b (-b5=a -p=n); individual nucleotide alignments were concatenated into a multi-gene alignment with partition information corresponding to SCOGs. Phylogenetic relationships between the four species were reconstructed using ten independent runs of IQ-TREE2 v.2.3.1 (- m MFP -bb 1000 -alrt 1000 -abayes -bnni) (Minh *et al*. 2020) on the partitioned alignment (concat tree hereafter). Ancestral nodes in the concatenated, partition tree were dated using IQ-TREE2 with the minimum calibration point: [*Gossypioides* + *Kokia*] ∼5.3 mya (Grover *et al*. 2017).

Concordance among individual gene trees was characterized for a subset of filtered genes as follows. We first removed single-gene alignments without parsimony-informative sites as estimated by IQ-TREE2 (-m MFP -n 0 -alninfo). We produced individual-gene trees for the remaining alignments (sg tree hereafter) using IQ-TREE2 (-B 1000 -m MFP). Finally, a concordance analysis was conducted with IQ-TREE2 (-t species.tree --gcf loci.treefile --prefix concordg) to compare concat and sg trees. We used PhyloPart (https://sourceforge.net/projects/phylopart/) and PhypartsPieCharts from the phyloscripts project (https://github.com/mossmatters/phyloscripts/tree/master) to visualize concordance between concat and sg trees, after collapsing nodes with low bootstraps support (BS<70).

We calculated pairwise d*N*/d*S* values for each SCOGs with CODEML (PAML v.4.10.7) under the basic model (model = 0; NSsites = 0) and the FmutSel codon fitness (CodonFreq = 7). We estimated median values and plotted pairwise d*N*, d*S*, and d*N*/d*S* values into density curves and boxplots using the dplyr v.1.1.4, gridextra v.2.3, and tidyverse v.1.3.2 modules in R.

The median of the d*S* distribution for the set of SCOGs analyzed above was used to estimate the synonymous substitutions rate per site per year (r) following the equation 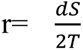 (https://ngdc.cncb.ac.cn/biocode/tools/BT000001/manual), where T is the estimated divergence time (5.3 mya) between *Gossypioides* and *Kokia* (Grover *et al*. 2017). We then used the average r for all *Kokia*-*Gossypioides* estimates to calculate divergence times within *Kokia* using the same formula.

## Results and Discussion

### Genome assembly and annotation

We report high-quality chromosome level genome assemblies for *Kokia cookei*, *K. drynarioides*, and *K. kauaiensis*. Chromosomes ranged 35.8-62.0 Mbp in size with a small amount of unresolved sequence (65,300-88,900 of Ns) per species (Table 1). Benchmarking Universal Single-Copy Ortholog (BUSCO) analysis revealed a high level (96.5-98.8 %) of completeness of the genome assemblies, with only 0.3-0.6 % fragmented and 0.8-3.0 % missing (Table 1). Our improved assembly for *K. drynarioide*s yielded 1,654 scaffolds (N50 = 40.5 Mbp) resulting in a total size of 552.4 Mbp, 512 Mbp of which assembled into twelve chromosomes. By comparison, the previous iteration of this genome consisted of 19,146 scaffolds (N50 = 176.7 kbp) amounting to 520.9 Mbp, and representing 95.6 % of genomic BUSCO groups (Grover *et al*. 2017).

**Table 1:** Assembly statistics and BUSCO scores for *Kokia cookei*, *K. drynarioides*, and *K. kauaiensis* genomes.

|  | *K. cookei* | *K. drynarioides* | *K. kauaiensis* |
| --- | --- | --- | --- |
| Scaffolds | 1,109 | 1,654 | 910 |
| N50 (Mbp) | 42.2 | 40.5 | 41.1 |
| Assembly length (bp) | 561,147,504 | 552,362,197 | 555,992,597 |
| Total length of Ns* | 88,900 | 76,800 | 65,300 |
| Repeats (%) | 63.95 | 64.81 | 63.69 |
| Complete BUSCO |  |  |  |
| Total | 96.50 % | 98.60 % | 98.80 % |
| Single | 84.40 % | 89.60 % | 87.80 % |
| Duplicated | 12.10 % | 9.00 % | 11.00 % |
| Incomplete BUSCO |  |  |  |
| Fragmented | 0.50 % | 0.60 % | 0.30 % |
| Missing | 3.00 % | 0.80 % | 0.90 % |

\* gaps in assembly filled with runs of 100 Ns

We used BRAKER3 to *de novo* annotate the three genomes, recovering 38,042 to 39,268 gene models per species (Table 2). BUSCO analysis of the annotation similarly resulted in 93.5-96.2 % complete orthologs (77.5-81.3 % single, 14.1-16.0 % duplicated) with 1.7-2.1% fragmented and only 2.1-4.6 % missing. We further assessed our annotations with orthology analyses using both Genespace and Orthofinder. When restricting the analyses to the three *Kokia* species, OrthoFinder (OF) found that the majority (97.5-97.8 %) of predicted genes fell into 32,480 orthogroups (OGs). Our analyses recovered 27,332 OGs containing all three species, 20,630 of these being single copy orthologs (SCOGs). We also found 486 species-specific OGs containing 601-818 genes per species (1.5-2.0 % of genes per species). *Kokia cookei* contains the largest portion of genes (818 genes, 2.0 %) in species-specific OGs (SSOGs). When adding *G. kirkii* into the analysis, OF identified 635-400 (1-1.6 %) and 1616 (3.9 %) SSOGs in *Kokia* and *Gossypioides*, respectively. By comparison, previous analysis found 5,188 and 4,400 unique genes in *G. kirkii* and *K. drynarioides*, respectively (Grover *et al*. 2017). *Kokia*-specific GENESPACE run organized the genes into 36,601 syntenic ortholog groups (SynOGs) containing genes from all three species, representing 15,971 genes more than what was found by OF. *Kokia* genomes contain a median of three exons per gene (125 bp in size) interleaved with introns ranging from 136-139 bp in size, the latter values slightly below what has been estimated for land plants (Figure S1; (Wu *et al*. 2013)). We also predicted 8133, 7339, and 8070 single-exon genes in *K. cookei*, *K. drynarioides* and *K. kauaiensis*, respectively, corresponding to 19-21 % of annotated protein coding genes. Interestingly, the sister clade *Gossypium* displays a median of four exons per gene, suggesting that the lineage leading to *Kokia* experienced genome-wide intron loss. While plants generally have smaller genes than other eukaryotes, mainly due to fewer and smaller exons per gene (Ramírez-Sánchez *et al*. 2016), it appears that *Kokia* genes may have traveled further in the trajectory of gene reduction compared to closely-related taxa; however, a broader generic sampling is required to phylogenetically characterize this intron-loss phenomenon.

**Table 2:** Gene statistics and relationships between *Kokia cookei*, *K. drynarioides*, and *K. kauaiensis* genomes.

|  | *K. cookei* | *K. drynarioides* | *K. kauaiensis* |
| --- | --- | --- | --- |
| Number of genes | 39268 | 38042 | 39242 |
| Complete BUSCO |  |  |  |
| Total | 93.30 % | 95.90 % | 96.1 % |
| Single | 77.7 % | 81.4 % | 81.1 % |
| Duplicated | 15.9 % | 14.0 % | 15.0 % |
| Incomplete BUSCO |  |  |  |
| Fragmented | 1.9 % | 2.0 % | 1.7 % |
| Missing | 4.5 % | 2.6 % | 2.2 % |
| OrthoFinder* |  |  |  |
| Genes in orthogroups (%) | 39076 (97.8) | 38323 (97.5) | 39101 (97.8) |
| Unassigned genes (%) | 880 (2.2) | 993 (2.5) | 875 (2.2) |

\* only non-overlapping genes on chromosome assemblies are considered

### Genomics and evolution of Kokia

We built a riparian plot for the *Kokia* genomes using *G. kirkii* as a reference, which shows the high syntenic stability within *Kokia* (Figure 2). In general, gene order is conserved between *K. drynarioides* and *K. kauaiensis*, with slight gene reshuffling in *K. cookei* and, interestingly, two major intra-chromosomal inversions (chromosomes 6 and 12) compared to *G. kirkii* (Figure 2) that were not previously described. Such genome conservation is also observed in the sister clade *Gossypium* (Chen *et al*. 2020). Given that our Genespace analyses were limited to sequences that were assembled into chromosomes, we observed a ∼600 kbp segment (∼ 660 genes) missing on chromosome 11 of *K. cookei* compared to the other genomes (Figure 2). This region is also present in *K. cookei* but could not be assembled with the rest of the genome with confidence.

**Figure 2:**
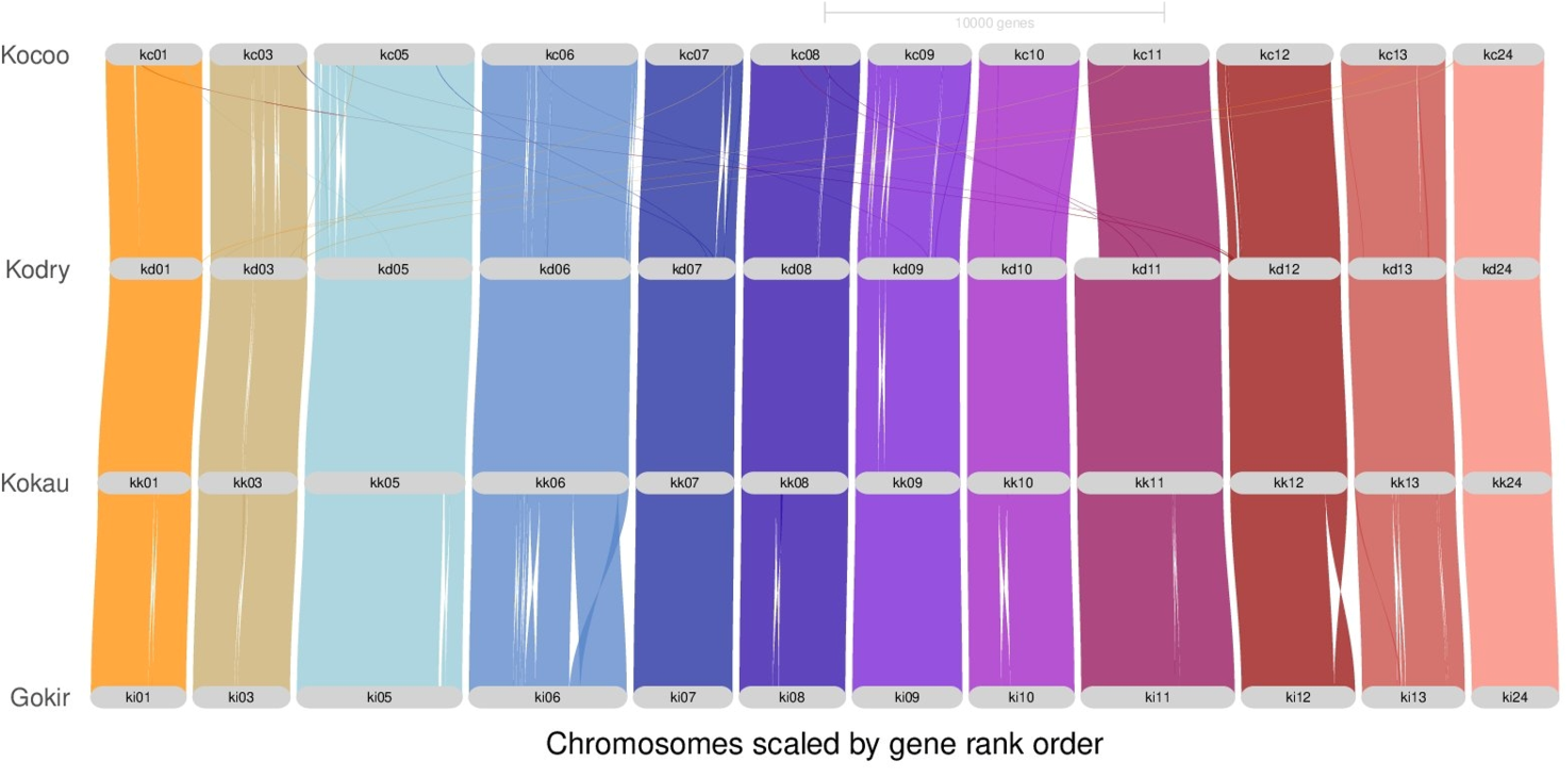
Gene order graph for the three *Kokia* species (n=12) sequenced in this study. Synteny, illustrated by coloured lines between chromosomes, was reconstructed with GENESPACE v.1.2.3. Chromosomes are numbered according to the reference, where chromosome 24 corresponds to the fused chromosomes 2 and 4 (Udall *et al*. 2019). Kocoo/kc: *K. cookei*; Kodry/kd: *K. drynarioides*; Kokau/kk: *K. kauaiensis*; Gokir/ki: *Gossypioides kirkii*.

We used RepeatMasker to identify 63.7-64.8 % of repeated sequences in the *Kokia* genomes, about half (32.05-33.68 %) classified as retroelements, and 26.73-27.72 % unclassified. As expected, *Gypsy/DIRS1* constituted the majority (20.2-21.3 %) of the identified repeats, followed by *Ty1/Copia* (5.4-7.1 %). These values are higher than previously described for *K. drynarioide*s (Grover *et al*. 2017), possibly due to the greater genome contiguity. Overall, the repeat landscape is highly conserved within the clade [*Kokia* + *Gossypioides*], whereas substantial gain and loss of repeats (32-63 %) have been described in the genome of members of the sister clade *Gossypium* (Grover *et al*. 2021) whose members also exhibit greater genome size variation (Hendrix and Stewart 2005). The high percentage of unclassified repeats found in both *Kokia* and *Gossypioides* (71-75.22 % of total interspersed repeats) suggest a hidden diversity of selfish elements that requires further investigation, notably by exploring the genome from the remaining taxon from that clade, i.e., *G. brevilanatum* and other members of tribe *Gossypieae*.

### Origin and diversification of Kokia

We reconstructed the phylogenetic relationship between the three *Kokia* species using *Gossypioides kirkii* as an outgroup. The Maximum Likelihood tree based on a multi-gene nucleotide alignment of 17,224 SCOGs (concat) containing 23,670 parsimony informative sites recovered the clade [*K. cookei* + *K. drynarioides*] sister to *K. kauaiensis* with maximum support (Figure 3).

**Figure 3:**
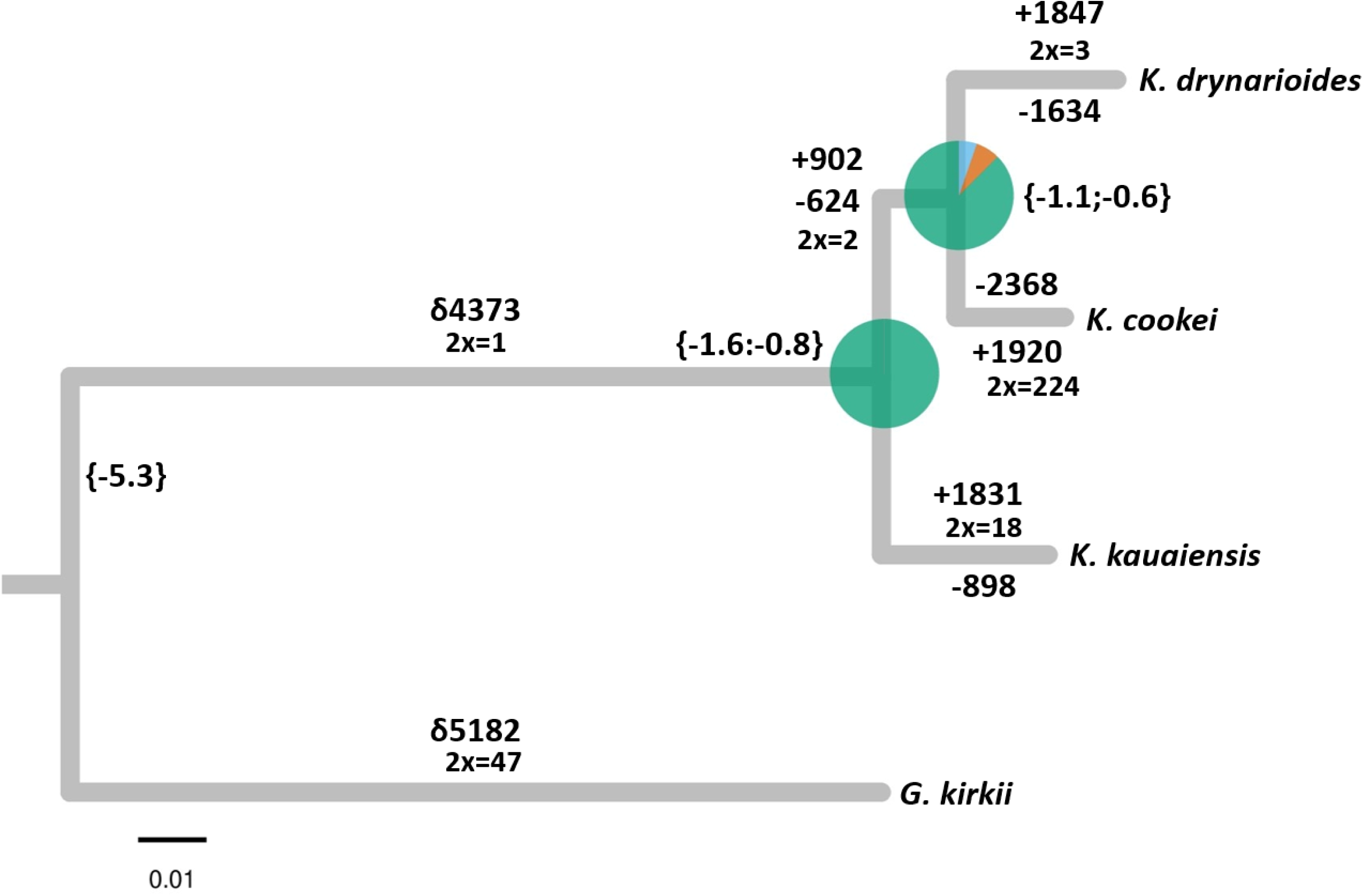
Dated phylogenetic relationships between the three *Kokia* species sequenced in this study using *Gossypioides kirkii* as outgroup. All nodes have 100 % bootstrap support. The tree was built with IQ-TREE2 using a partitioned multiple gene alignment of 17,224 single copy orthologs reconstructed by OrthoFinder containing 23,897 parsimony informative sites. Dating was performed with IQ-TREE2 using minimum calibration points as follows: [*Gossypioides* + *Kokia*] ∼5.3 mya; [*K. cookei* + *K. drynarioides* + *K. kauaiensis*] =1.2 mya, predicting the split between [*K. cookei* + *K. drynarioides*] around 0.8 mya. Branches also display differences (δ), gain (+) and loss (-) of orthogroups identified by GENESPACE; 2x represents the number of OGs that experienced a two-factor expansion in a given branch of the tree. Piecharts reflect the concordance of 8,973 single-gene trees to the concatenated tree, obtained with PhyloPart and PhypartsPieCharts after collapsing nodes with low bootstraps support (BS<70), where green corresponds to recovered node, blue to main alternative node, and orange to other alternative relationships.

We also produced phylogenetic trees for 8,973 single-gene (sg) alignments that contained parsimony informative sites (PIS) and displayed no saturation according to the Xia *et al*. saturation test implemented in DAMBE v.7.3.32 (Xia *et al*. 2003, 2009). Most sg trees agree with the relationships obtained in the concat tree (Figure 3). Interestingly, 644 sg trees preferred the alternative [*K. cookei* + *K. kauaiensis*] sister to *K. drynarioides*, whilst 473 sg trees supported [*K. drynarioides* + *K. kauaiensis*] sister to *K. cookei*. These numbers dropped to 393 and 288, respectively, when removing nodes with bootstrap support <70.

Divergence among *Kokia* species is relatively recent (∼1.2 mya) compared to the divergence between *Kokia* and *Gossypioides* (∼5.3 mya), although this estimate includes only extant species (e.g., without *K. lanceolata* and any possible species endemic to now submerged islands). We found the subsequent divergence of the [*K. cookei* + *K. drynarioides*] clade to be approximately 0.8 mya when using −5.3 as minimum calibration point (Grover *et al*. 2017). This is congruent with the fact that *Kokia cookei* and *K. drynarioides* were originally described from the younger major islands of the Hawaiian archipelago, namely Moloka‘i and Hawai‘i, respectively, while the more distantly related *K. kauaiensis* inhabits mesic forests of the older Kaua‘i Island. Notably, the divergence time between *Gossypioides* and *Kokia* is older than the estimated emergence of the more recent Hawaiian islands, namely 4.7 to 0.5 mya (Price and Clague 2002). Consequently, the ancestor of *Kokia* likely first colonized the archipelago starting with the lower older islands, including some that are now under the sea level, as proposed by an earlier study (Morden and Yorkston 2018). Further diversification of the genus came along with the emergence of the younger volcanic islands, a pattern similar to other biota of the Hawaiian Archipelago (Price and Clague 2002).

We estimated pairwise genetic divergence among the three *Kokia* species for both synonymous (d*S*) and nonsynonymous (d*N*) substitutions using the 17,224 filtered SCOGs that exclude those with d*S* > 1 (Table 3). Median values for d*S* and d*N* within *Kokia* were low, similar to values estimated for tetraploid cotton species, which radiated approximately 1-2 mya after undergoing a severe polyploid bottleneck (Chen *et al*. 2020). While the mean d*N*/d*S* values were similar for *K. cookei* versus *K. kauaiensis* and *K. drynarioides* versus *K. kauaiensis* (0.3842; Figure S2), d*S* and d*N* were both lower for the sister species comparison, *K. drynarioides* versus *K. cookei*. Pairwise comparisons for d*N* and d*S* values between each of the three *Kokia* species versus *Gossypioides kirkii* were similar, for all comparisons (d*N* ∼ 0.011, d*S* ∼ 0.020; Table 3). Interestingly, the median d*S* value for *K. cookei* versus *K. drynarioides* was less than the median for d*N* (Table 3), possibly reflecting the stochasticity inherent in small numbers of substitutions and possibly some unknown targets of selection that differ between species.

**Table 3:**
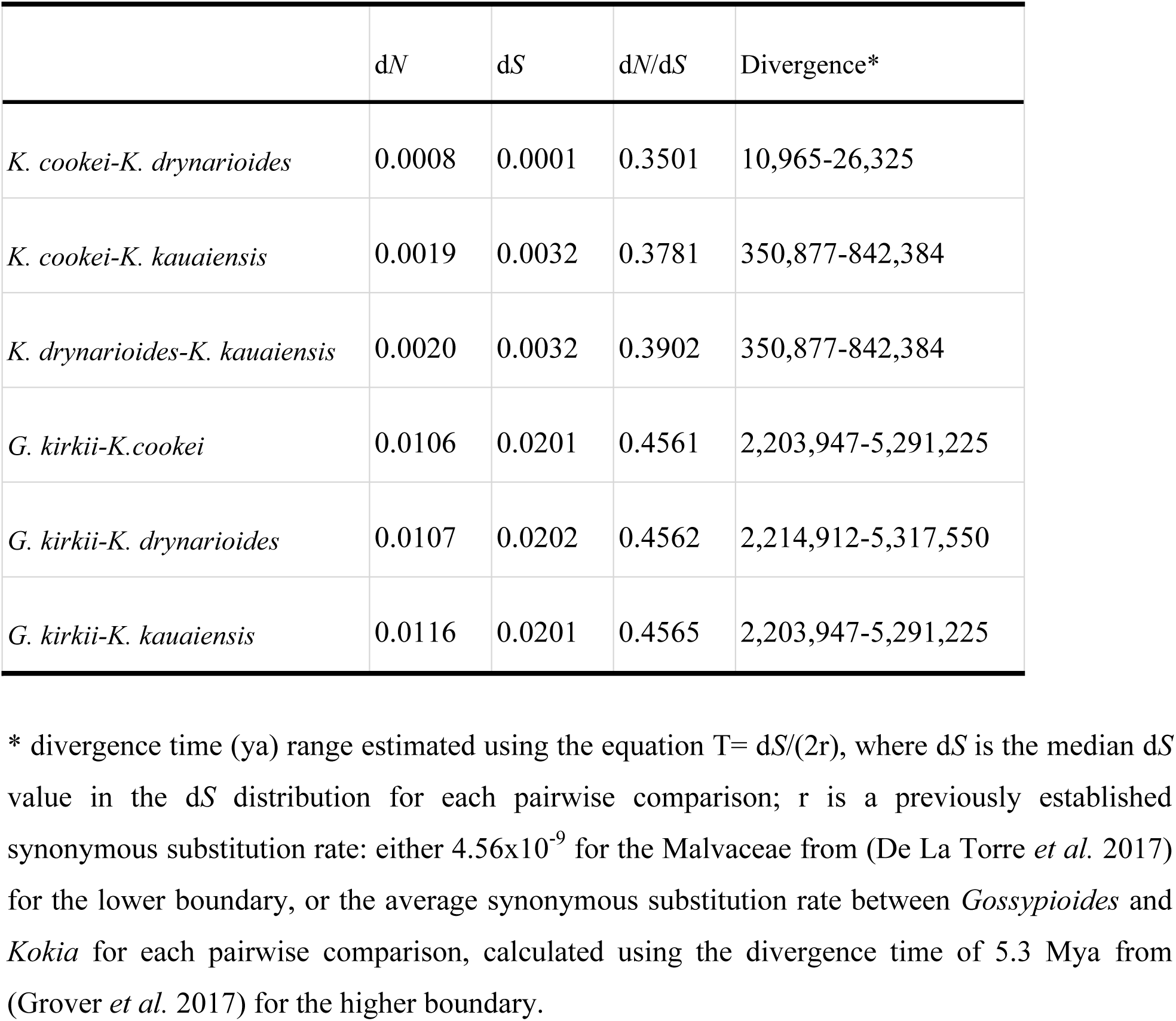
Median pairwise genetic divergence between *Kokia cookei*, *K. drynarioides*, *K. kauaiensis,* and *Gossypioides kirkii* genomes.

Previous research estimated the divergence time between *Kokia* and *Gossypioides* to be ∼5.3 mya. Using this approximate divergence time, we observe the mutation rate (Nei and Kumar 2000) between *Kokia* and *Gossypioides* (r) to average 1.9 x10^-9^ substitutions per site per year, lower than original estimated rate of 4.56×10^-9^ substitutions/site/year for Malvaceae (De La Torre *et al*. 2017) used originally in the previous comparison of *K. drynarioides* and *G. kirkii* (Grover *et al*. 2017). The latter rate was estimated based on 13,643 SCOGs using the more distantly related *Gossypium raimondii*, which could explain an overestimation of the substitution rate.

Using the newly calculated average rate of 1.9 x10^-9^ substitution per site per year, we estimated a divergence time of ∼842,000 ya between [*K. cookei* + *K. drynarioides*] and *K. kauaiensis* (median d*S* 0.003), and a mere ∼26,000 ya between *K. cookei* and *K. drynarioides* (Table 3). We observed that the d*S* method estimated divergence time ranges (350,877-842,384 years for [*K. cookei*/*K. drynarioides*-*K. kauaiensis*] and 10,965-26,325 years for [*K. cookei*-*K. drynarioides*]) are lower than those estimated by the phylogenetic analysis (Figure 3; Table 3). Estimating divergence time using substitution values can be problematic when high variance in the distribution of d*S* (and d*N*) are observed across genes (Figure 4; Figure S2).

**Figure 4:**
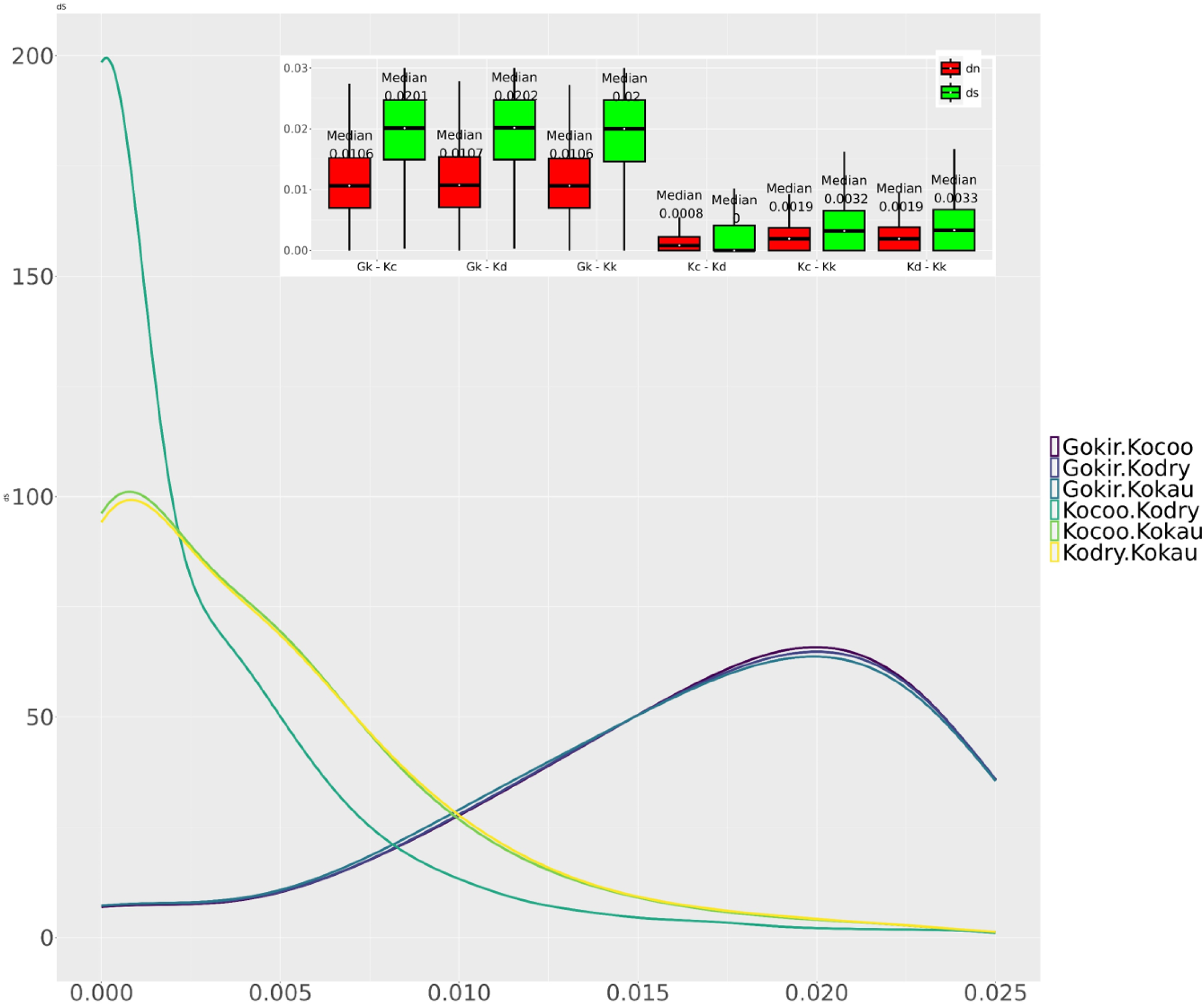
Distribution of substitution rates between pairwise comparisons of the three *Kokia* genomes reported here and *Gossypioides kirkii*. The curve represents the frequency distribution of pairwise d*S* comparisons calculated for 17,224 single copy orthologs (identified by OrthoFinder v.2.5.4) with CODEML (PAML v.4.10.7) under the basic model (model = 0; NSsites = 0), after removing those with d*S* >1. Inset contains box plots of both synonymous (red) and nonsynonymous (green) substitution values (including the median) for each pairwise comparison for the same gene set. Gokir/Gk: *Gossypioides kirkii*; Kocoo/Kc: *Kokia cookei*; Kodry/Kd: *K. drynarioides*; Kokau/Kk: *K. kauaiensis*.

### Gene content evolution in Kokia

Previous research (Grover *et al*. 2017) suggested a disproportionate rate of gene loss versus gain in *Kokia* (4:1, respectively), noting that this study was based on a single representative of *Kokia* (i.e., *K. drynarioides*), resulting in a difference in gene content between the two genera totaling 6,486 genes. Here, we expanded the investigation of gene loss and gain in *Kokia* using the GENESPACE classifications of orthogroups for the four species (i.e., three *Kokia* and the outgroup *Gossypioides kirkii*). We found 5,182 genes in the lineage leading to *Gossypioides* that are absent in *Kokia* (Figure 3), representing either gains in the *Gossypioides* lineage or losses in the *Kokia* lineage. By comparison, only 70 % as many genes (δ4,373; Figure 3) were absent in *Gossypioides* but present either in [*K. cookei* + *K. kauaiensis*] or in all three *Kokia* species, which would indicate gains in the lineage leading to *Kokia* or losses in *Gossypioides*. Although the present study does not distinguish between gains and losses, it is notable that more genes unique to *G. kirkii* versus to *Kokia* have been identified than previously estimated (Grover *et al*. 2017), possibly due to the inclusion of multiple *Kokia* representatives. Whereas the previous estimate suggested 3,747 genes unique to *Kokia* and 2,739 unique to *Gossypioides* (1.4:1 unique *Kokia* to unique *Gossypioides*), the current estimate suggests a ratio closer to 0.7:1 between any *Kokia* and *G. kirkii*, a substantial difference likely resulting from both increased in taxa sampling and the inclusion of synteny in orthology search. When looking at copy number variation (CNV) within gene families, we identified the largest expansion occurred in *K. cookei*, where 224 orthogroups (OGs) underwent two-fold expansions. We found two-fold expansion in 3, 18, and 47 OGs in *K. drynarioides*, *K. kauaiensis*, and *G. kirkii*, respectively (Figure 3). Further exploration of the genome of *G. brevilanatum* will be helpful in understanding gene family contraction and extension in the [*Kokia* + *Gossypioides*] clade.

Within *Kokia*, gene loss and gain were variable among lineages, with the fewest changes in the lineage leading to *K. kauaiensis* and the greatest in the lineage leading to the grafted species, *K. cookei*. The previously sequenced *K. drynarioides* was intermediate between the other two species and exhibited nearly balanced numbers of gains and losses (1,847 gains versus 1,634 losses). In contrast, *K. kauaiensis* exhibited more than twice as many gains (1,831) as losses (898), whereas *K. cookei* exhibited the opposite (1,920 gains versus 2,368 losses). An additional 902 gains and 624 losses have occurred in the past ∼1.2 million years in the [*K. drynarioides* + *K. cookei*] clade compared to *K. kauaiensis*. Notably, these observations differ somewhat from a previous study that estimated twice as many losses than gains in *G. kirkii* and *K. drynarioides*; however, these observations were based solely on OrthoFinder analyses and a lower quality genome sequence (Grover *et al*. 2017). Our study shows how including additional high-quality genomes and improving genome sequences (e.g., *K. drynarioides*) can permit a more thorough understanding of evolutionary change within and among genera (here, *Kokia*).

## Conclusion

*Kokia* is an endangered genus of insular plants endemic to Hawai‘i comprising three extant species. The reference genomes generated for these three species represent an important step toward understanding the genetic makeup of *Kokia* species, thereby facilitating conservation efforts (Theissinger *et al*. 2023). These resources form the foundation for future resequencing efforts that will provide insight into population structure and processes within each *Kokia* species. This in turn will provide data regarding genetic diversity, mutation loads, and relatedness information to inform conservation efforts aimed at preserving the diversity and population viability of this endangered genus (Werden *et al*. 2020; Pegueroles *et al*. 2024). Additionally, differences in gene content and their functional significance require further study. Finally, these resources will serve as a reference for exploring the only available herbarium specimen of the extinct *K. lanceolata* collected in late 19^th^ century, thereby contributing to our understanding of the evolution of island endemic genera such as *Kokia* and their survivability in the face of human disturbances.

## Data availability

The assembled *Kokia* genome sequences are available at NCBI under BioProject: PRJNA1087748 and CottonGen (https://www.cottongen.org/). Raw sequencing reads are also available at NCBI SRA under BioProject: PRJNA1087748. Relevant code is available at https://github.com/Wendellab/ThreeKokiaGenomes.

## Acknowledgements

We thank Waimea Valley (arboretum and botanical garden in Haleiwa, HI, USA) and Plant Collections Specialist David Orr for providing fresh *Kokia* tissue and records regarding these specimens. We thank the USDA-ARS (58-6066-0-066, Genomics of Malvaceae) for their financial support. This work was made possible by the USDA Supercomputer Atlas funded through the SCINet initiative. We thank the Iowa State University ResearchIT unit for computational resources and support. EK would like to extend his thanks to Weixuan Ning and OpenAI’s ChatGPT for their help in coding.

